## Supplementary material for "Translation in Bacillus subtilis is spatially and temporally coordinated during sporulation": Document S1

```
A
RpsN      MAKKSMIAKQQ--------------------------RTP--KFKVQEYTRCERCGRPHSVIRKFKLCRICFRELAYKGQIPGVKKASW
RpsNB     MAKKSKVAKELKRQQLVEQYAGIRRELKEKGDYEALSKLPRDSAPGRLHNRCMVTGRPRAYMRKFKMSRIAFRELAHKGQIPGVKKASW
          ***** :**:                           : *  .   : :.**   ***:: :****:.**.*****:************

B
RpmE      MKAGIHPNFKKATV---------------KCACGNEFETGSVKEEVRVEICSECHPFYTGRQKFASADGRVDRFNKKYGLK-
RpmEB     MKEGIHPKNHKVIFQDVNSGYRFLSTSTKTSNETAEWEDGNTYPVIKVEVSSDTHPFYTGRQKFNEKGGRVEQFKKRYNMGK
          ** ****: :*. .               ..    *:* *..   ::**:.*: ********** . .***::*:*:*.:

C
RpmGA     MRVNITLACTECGERNYISKKNKRNNPDRVEFKKYCPRDKKSTLHRETK
RpmGC     MRVNVTLACTETGDRNYITTKNKRTNPDRLELKKYSPRLKKYTLHRETK
```

**Supplemental Figure S1.** Alignment of paralogs of S14, L31, and L33 ribosomal proteins, respectively **(A)** RpsNB, **(B)** RpmEB, **(C)** RpmGC. The CXXC zinc-binding motif is marked in bold.

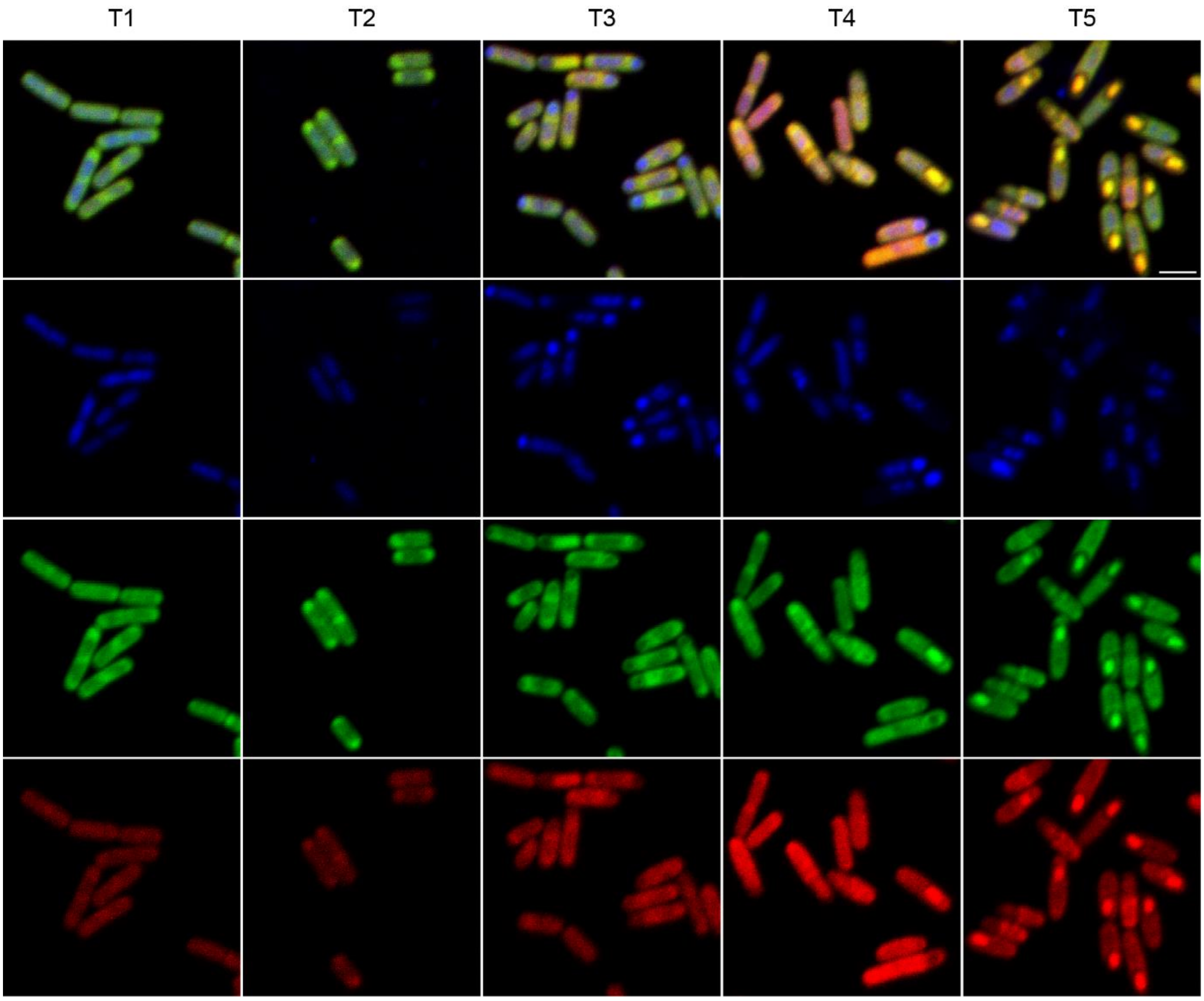


**Supplemental Figure S2.** The ribosomal localization of RpmEB at 1, 2, 3, 4, and 5 hours post-sporulation induction. Cells were stained with DAPI (blue) to visualise the chromosome. The ribosomal proteins were tagged with fluorescent protein tags – RpsB-GFP (green) and RpmEB-mCherry (red). Scale bar is 2µm.

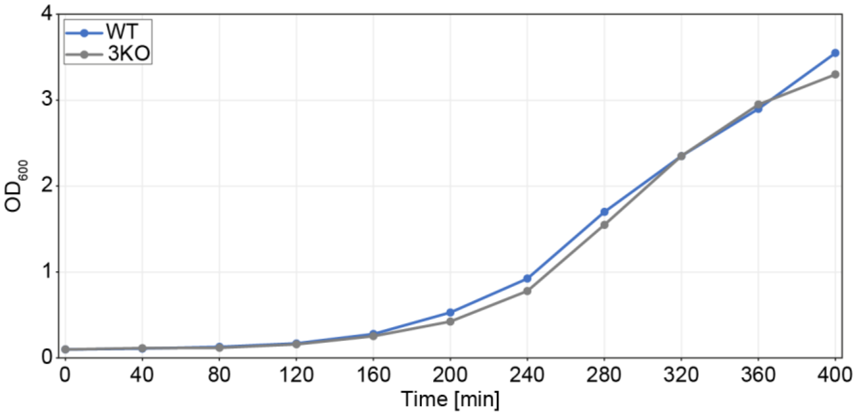


**Supplemental Figure S3.** Growth Curves in CH Medium at 37°C for WT (blue) and 3KO (gray) Strains.

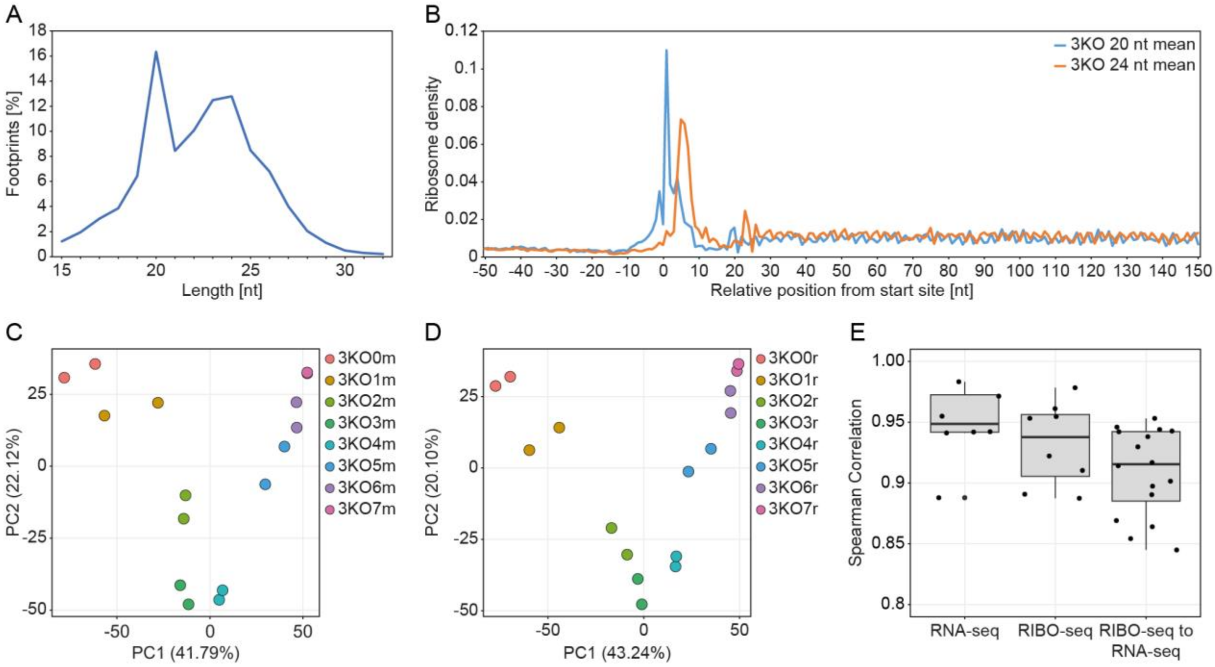


**Supplemental Figure S4.** Quality Control of 3KO Data. **(A)** Length distribution of ribosomal footprints (RPFs) from all timepoints with duplicates. **(B)** Metagene plots of 3' assigned RPFs on ORF for 20 (blue) and 24 nt long (orange) footprints. **(C)** and **(D)** Principal component analysis plots of duplicate samples of the transcriptome (RNA-seq) and translatome (Ribo-seq) data for sporulating *Bacillus subtilis* at different timepoints: 0 to 7 hours post sporulation induction. **(E)** Distribution of Spearman's rho values of correlation between duplicates for RNA-seq and Ribo-seq and between transcriptome (RNA-seq) and translatome (Ribo-seq) dataset pairs.

**Supplemental Table S4. (A)** The sporulation/germination efficiency was measured as the ratio of CFUs resulting from fully sporulated cultures before and after heat treatment (40 min at 90°C). **(B)** Sporulation efficiency was calculated as the ratio between cells with asymmetric septum and cells without, for both WT and 3KO strains (n > 600), at 2, 3, and 4 hours post-sporulation induction.

| **A. Sporulation/germination efficiency before and after heat treatment** | | | |
|---|---|---|---|
| **Strain** | **$10^{-5}$** | **$10^{-5}$ heated** | **%** |
| **WT** | 160 | 101 | 63.1% |
| | 125 | 77.3 | 61.9% |
| | 142.3 | 83.3 | 58.5% |
| | 181.3 | 119 | 65.5% |
| | 160 | 94 | 58.7% |
| **3KO** | 212 | 37 | 17.4% |
| | 237 | 53.3 | 22.5% |
| | 161.7 | 71.7 | 44.3% |
| | 244 | 106.3 | 23% |
| | 267.6 | 61.6 | 43.6% |
| **B. Sporulation efficiency at 2, 3, and 4 hours post-sporulation induction** | | | |
| **Strain** | **T2** | **T3** | **T4** |
| **WT** | 370/870 = 42% | 392/505 = 77% | 569/730 = 78% |
| **3KO** | 171/464 =37% | 318/487 = 65% | 415/518 = 80% |

**Supplemental Table S6.** Lists of **(A)** strains, **(B)** primers, **(C)** plasmids used in this study, and **(D)** adaptors and primers used to prepare cDNA libraries (from Multiplex Small RNA Library Prep Set for Illumina NEB #E7300 [1]).

| **A. Strains** | | |
|---|---|---|
| **Strain** | **Genetic description** | **Reference** |
| **WT** | *Bacillus subtilis* 168 | Burkholder and Giles, 1947 [2]; Spizizen, 1958 [3] |
| **Single KO** | B. subtilis 168 Δ*rpmEB* $Kan^R$ | This study |
| **2KO** | B. subtilis 168 Δ*rpmEB* Δ*rpsNB* $Kan^R$ $Cm^R$ | This study |
| **3KO** | B. subtilis 168 Δ*rpmEB* Δ*rpsNB* ΔrpmGC $Kan^R$ $Cm^R$ $Ery^R$ | This study |
| **WT-GFP** | *Bacillus subtilis* 168 *rpsB*:*gfp* $Sp^R$ | This study |
| **3KO-GFP** | B. subtilis 168 Δ*rpmEB* Δ*rpsNB* ΔrpmGC *rpsB*:*gfp* $Kan^R$ $Cm^R$ $Ery^R$ $Sp^R$ | This study |

| **B. Primers** | |
|---|---|
| **Primer** | **Sequence** |
| **UPFOR_rpmEB_1** | AGGAAGCACCAAAATTAAAATAGCAGG |
| **UPREV_rpmEB_1** | CACTGCCCGCTTTCCAGTCGGGGGGTATCTCCTTTCAATAAATCG |
| **MIDFOR_rpmEB_2** | CGATTTATTGAAAGGAGATACCCCCCGACTGGAAAGCGGGCAGTG |

| | |
|---|---|
| **MIDREV_rpmEB_2** | GACAAACCCTCAGGCCTGCCGTTATCGACAGCGGAATTGACTC |
| **DOWNFOR_rpmEB_3** | GAGTCAATTCCGCTGTCGATAACGGCAGGCCTGAGGGTTTGTC |
| **DOWNREV_rpmEB_3** | GCAGCATCGGCGTCTCCGACTTGGAC |
| **UPFOR_rpsNBcm_1** | ATTATCATTTTTGCAGTGGTTGGAAAGCTG |
| **UPREV_rpsNBcm_1** | CTTTATTATACAGATCTCCATGTCACATAGCCTCCCTTTAAATCG |
| **MIDFOR_rpsNBcm_2** | CGATTTAAAGGGAGGCTATGTGACATGGAGATCTGTATAATAAAG |
| **MIDREV_rpsNBcm_2** | CGGCGCGCTGTAATCGGCCGGAGTTTTTTCCACAAGAGGACGCTTTATTCTTCC |
| **DOWNFOR_rpsNBcm_3** | GGAAGAATAAAGCGTCCTCTTGTGGAAAAAACTCCGGCCGATTACAGCGCGCCG |
| **DOWNREV_rpsNBcm_3** | TCATTCTGGAAGGATATGCGAGAGCAATTG |
| **UPFOR_rpmGCe_1** | GGCGGCTCCATTGCAAGAAGGGCAACCATCTG |
| **UPREV_rpmGCe_1** | CCGTCTTATCTCCCATTATATCTTTTTTTTATTATACAACGTCATCACAAATTG |
| **MIDFOR_rpmGCe_2** | CAATTTGTGATGACGTTGTATAATAAAAAAAAGATATAATGGGAGATAAGACGG |
| **MIDREV_rpmGCe_2** | CGTAAAAAAAGACCGGGCCGTAAGGGAGTAGTATACCTAATAATTTATCTAC |
| **DOWNFOR_rpmGCe_3** | GTAGATAAATTATTAGGTATACTACTCCCTTACGGCCCGGTCTTTTTTTACG |
| **DOWNREV_rpmGCe_3** | GCGGTTCATTTGAGCAGTTTATATGACGGGAAAGAAATGACTTG |
| **UPFOR_rpsB_1** | GATACCTACGCCTCGTTTAGAATTCGCGGCGCAATC |
| **UPREV_rpsB_1** | CATTGATCCGCTGCCTGATCCGGACGCAGTTGTTGTTTCTGTTTC |
| **MIDFOR_rpsB-GFP_2** | GAAACAGAAACAACAACTGCGTCCGGATCAGGCAGCGGATCAATG |
| **MIDREV_rpsB-GFP_2** | GTATTTTTCCGTTAATCAAATTGCTCATTCACTTATAGAGTTCATCCATACC |
| **MIDFOR_rpsBsp_3** | GGTATGGATGAACTCTATAAGTGAATGAGCAATTTGATTAACGGAAAAATAC |
| **MIDREV_rpsBsp_3** | GTCCCTCTTATCACCTTTTGAATAGGTAATTGAGAGAAGTTTCTATAGAATTTTTC |
| **DOWNFOR_rpsB_4** | GAAAAATTCTATAGAAACTTCTCTCAATTACCTATTCAAAAGGTGATAAGAGGGAC |
| **DOWNREV_rpsB_4** | GTGTCTGCGCTCCGTAATATTCAACCGTTACTTTATCTAATAATG |
| **C. Plasmids** | |
| **Name** | **Description** |
| **pAPNC-kan** | Used as a template for kanamycin resistance cassette |
| **pAPNC-erm** | Used as a template for erythromycin resistance cassette |
| **pAPNC-cm** | Used as a template for chloramphenicol resistance cassette |
| **pSHP2** | Used as a template for GFP and spectinomycin resistance cassette |
| **D. Adaptors and primers used to prepare cDNA libraries** | |
| **Primer** | **Sequence** |
| **NEBNext SR Primer for Illumina** | AATGATACGGCGACCACCGAGATCTACACGTTCAGAGTTCTACAGTCCG*A |
| **NEBNext SR RT Primer for Illumina** | AGACGTGTGCTCTTCCGATCT |
| **NEBNext Index 1 Primer for Illumina** | CAAGCAGAAGACGGCATACGAGATCGTGATGTGACTGGAGTTCAGACGTGTGCTCTTCCGATC*T |
| **NEBNext Index 2 Primer for Illumina** | CAAGCAGAAGACGGCATACGAGATACATCGGTGACTGGAGTTCAGACGTGTGCTCTTCCGATC*T |
| **NEBNext Index 3** | CAAGCAGAAGACGGCATACGAGATGCCTAAGTGACTGGAGTTCAGACGTGTGCTCTT |

| | |
|---|---|
| **Primer for Illumina** | CCGATC*T |
| **NEBNext Index 4 Primer for Illumina** | CAAGCAGAAGACGGCATACGAGATTGGTCAGTGACTGGAGTTCAGACGTGTGCTCT TCCGATC*T |
| **NEBNext Index 5 Primer for Illumina** | CAAGCAGAAGACGGCATACGAGATCACTGTGTGACTGGAGTTCAGACGTGTGCTCTT CCGATC*T |
| **NEBNext Index 6 Primer for Illumina** | CAAGCAGAAGACGGCATACGAGATATTGGCGTGACTGGAGTTCAGACGTGTGCTCTT CCGATC*T |
| **NEBNext Index 7 Primer for Illumina** | CAAGCAGAAGACGGCATACGAGATGATCTGGTGACTGGAGTTCAGACGTGTGCTCTT CCGATC*T |
| **NEBNext Index 8 Primer for Illumina** | CAAGCAGAAGACGGCATACGAGATTCAAGTGTGACTGGAGTTCAGACGTGTGCTCTT CCGATC*T |
| **NEBNext Index 9 Primer for Illumina** | CAAGCAGAAGACGGCATACGAGATCTGATCGTGACTGGAGTTCAGACGTGTGCTCTT CCGATC*T |
| **NEBNext Index 10 Primer for Illumina** | CAAGCAGAAGACGGCATACGAGATAAGCTAGTGACTGGAGTTCAGACGTGTGCTCTT CCGATC*T |
| **NEBNext Index 11 Primer for Illumina** | CAAGCAGAAGACGGCATACGAGATGTAGCCGTGACTGGAGTTCAGACGTGTGCTCT TCCGATC*T |
| **NEBNext Index 12 Primer for Illumina** | CAAGCAGAAGACGGCATACGAGATTACAAGGTGACTGGAGTTCAGACGTGTGCTCT TCCGATC*T |
| **NEBNext 3´ SR Adaptor for Illumi** | rAppAGATCGGAAGAGCACACGTCT-$NH_2$ |
| **NEBNext 5´ SR Adaptor for Illumina** | rGrUrUrCrArGrArGrUrUrCrUrArCrArGrUrCrCrGrArCrGrArUrC |

Where * indicates phosphorothioate bond
